# Intestinal parasitosis, anaemia and risk factors among pre-school children in Tigray Region, Northern Ethiopia

**DOI:** 10.1101/744938

**Authors:** Araya Gebreyesus Wasihun, Mekonen Teferi, Letemichal Negash, Javier Marugán, Dejen Yemane, Kevin G. McGuigan, Ronan M. Conroy, Haftu Temesgen Abebe, Tsehaye Asmelash Dejene

**Affiliations:** College of Health Sciences, Department of Medical Microbiology and Immunology, Mekelle University, Tigray, Ethiopia; College of Natural and Computational Sciences, Department of Biology, Mekelle University, Tigray, Ethiopia; Department of Chemical and Environmental Technology, Universidad Rey Juan Carlos, C/Tulipán s/n, 28933 Móstoles, Madrid, Spain; College of Health Sciences, School of Public Health, Department of Environmental Health, Mekelle University, Tigray, Ethiopia; Dept. of Physiology and Medical Physics, Royal College of Surgeons in Ireland (RCSI), Dublin 2, Ireland; Data Science Centre, Royal College of Surgeons in Ireland (RCSI), Dublin 2, Ireland; College of Health Sciences, School of Public Health, Department of Bio Statistics, Mekelle University, Tigray, Ethiopia; College of Health Sciences, Department of Medical Microbiology, Aksum University, Tigray, Ethiopia

**Keywords:** Intestinal parasite, anaemia, pre-school age children, risk factors, Tigray

## Abstract

**Background:** Intestinal parasitic infections (IPIs) and anaemia are major health problems. This study assessed the prevalence of IPI, anaemia and associated factors among pre-school children in rural areas of the Tigray region, northern Ethiopia.

**Methodology/Principal Finding:** A community based cross-sectional study was conducted among 610 pre-school children in rural communities of Northern Ethiopia from June 2017 to August 2017. Stool specimens were examined for the presence of trophozoites, cysts, oocysts, and ova using direct, formal-ethyl acetate concentration, Kato–Katz, and Ziehl-Neelsen techniques. Haemoglobin was measured using a HemoCue spectrometer. Among the 610 pre-school children participated in the study, prevalence of IPIs and anaemia were 58% (95% CI: 54.1–61.9%) and 21·6% (95% CI: 18·5% to 25·1%), respectively. Single, double, and triple parasitic infections were seen in 249 (41%, 95% CI: 37% to 45%), 83 (14%, 95% CI: 11% to 17%), and 22 (3.6%, 95% CI: 2.4% to 5.4%) children, respectively. Of the seven intestinal parasitic organisms recorded from the participants, *Entamoeba histolytica/dispar* was the most prevalent 220 (36.1%) followed by *Giardia lamblia* 128 (20.1%), and *Hymenolepis nana* 102 (16.7%). Mixed infections were common among *G. lamblia, E. histolytica/dispar* and *Cryptosporidium* spp. oocyst. Age 48-59 months prevalence ratio (PR = 1·078, P=0·009), child deworming (PR= 1.2; 95% CI=1.00-1.4, p= 0.045), and having two or more children aged under five (PR=0.76, 95% CI= 0.61-0.95, p=0.015) were independent predictors for IPIs. Anaemia was associated with proper disposal of solid waste (PR= 1.5, 95% CI=0.1.1-2.10, p=0.009). Eating raw meat (PR=0.49, 95% CI=0.45-0.54, p=0.000), any maternal education (PR=0.64 95% CI=0.52-0.79, p=0.000), and household water treatment (PR=0.75, 95% CI=0.56-1.0, p=0.044) were associated with lower prevalence of anaemia.

**Conclusions:** More than half of the children were infected with intestinal parasites and one in five were anaemic. This study has identified a number of potentially modifiable risk factors to address the significant prevalence of IPIs and anaemia in these children. Improvements in sanitation, clean water, hand hygiene, maternal education could address both short and long-term consequences of these conditions in this vulnerable population.

**Author Summary:** Intestinal parasitic infection and anaemia are public health problems in developing counties. Children due to their immature immune systems and frequent exposure to unhygienic environments are at high risk which in turn put them at an increased risk of malnutrition and growth deficits. Similarly, childhood anaemia impairs physical growth, impairs immune function and weakens motor development. The finding of this study reveals more than half of the children were infected. *Entamoeba histolytica/dispar, Giardia lamblia* and *Hymenolepis nana* were dominant parasites. Multiple infections was common among *Giardia lamblia, Entamoeba histolytica/dispar* and *Cryptosporidium* spp. Oocyst. Children aged 48-59 months were more infected with intestinal parasites. Soli transmitted helminths in this study was low. 21.5% of the children were anaemic and was associated with disposal of solid waste and presence of domestic animals. However, Eating raw meat, maternal education and household water treatment were found preventive of anaemia in the study. It seems worth understanding the prevalence and effects of parasitic infection and anaemia in this vulnerable group to design an appropriate interventions. Finally, if parasite transmission and anaemia is to be significantly prevented control programs such as improving sanitation, clean water, maternal education may be critical in this vulnerable age groups.

## Introduction

Intestinal parasitic infections (IPIs) are an important cause of morbidity and mortality worldwide (1,2) affecting about 3.5 billion people globally (3). IPIs are endemic in resource-limited regions due to high population density, low access to improved water sources, low latrine availability, poor hygiene conditions, low health awareness, and limited economic resources (4,5). Helminths such as *Ascaris lumbricoides*, Hookworm, *Strongloide stercolaris* and *Trichuris trichiura*, and enteric protozoan parasites such as *Entamoeba histolytica, Giardia lamblia* and *Cryptosporidium* spp. cause high incidences of health problems especially in children in low to middle income countries (6–9).

Children due to their immature immune systems and frequent exposure to unhygienic environments are at high risk for IPI including helminths (10), and protozoa (11). These infections are common during the period of life most critical for physical and cognitive development, hence are linked with an increased risk of childhood malnutrition and growth deficits (12). Poor health in children also results in deficits in cognitive development and educational achievements (13–15).

As with IPIs, anaemia remains a public health problem affecting both developing and developed countries with major consequences for human health as well as social and economic development (16). Globally, 2011 data indicate that 43% of children under-five were anaemic, with a higher prevalence in the developing world, specifically South Asia and East Africa, being 58% and 55%, respectively (17). Sub-Saharan Africa is one of the most affected regions with 53.8% of children under-five suffering from anaemia (18). Causes of anaemia include folate and iron deficiencies (19,20), malaria (21–23), infections (e.g., intestinal helminths), and diarrhoea (23). Childhood anaemia has many irreversible impacts; it impairs physical growth (24,25), impairs immune function, increases susceptibility to infections (26,27), and weakens motor development leading to reduced cognitive ability (16,28–31), poor school performance (32), and short or long term mortality in acute severe cases (20).

The 2016 Demographic and Health Survey of Ethiopia (EDHS) report showed that the national prevalence of anaemia among children aged 6 to 59 months was 57% (33), which exceeds the 40% threshold set by the World Health Organization (WHO) classification of anaemia as a severe public health problem (18). The prevalence of anaemia in the Tigray regional state in the EDHS report 54% is marginally below the national average (33).

Most studies conducted in Ethiopia on the prevalence of IPIs are on school-age children. Studies conducted among pre-school age children have shown wide variance in IPI prevalence from 21.2 % to 85.1% (34–40). Similarly, national studies on anaemia prevalence report prevalence from 32% to 37.3 % (41–43). These studies, however, are focused on either soil transmitted helminths alone (34,35,38) or symptomatic hospitalised children (39), or investigated anaemia alone. The association of IPI and anaemia has not been well addressed. Furthermore, none of the above studies used modified Ziehl-Neelsen techniques to detect *Cryptosporidium* spp., the second most causative agent of diarrhoea among children under five next to rotavirus (44).

There is scarcity of data on the prevalence of IPI, anaemia, and associated risk factors among pre-school children in the study area. However, the prevalence is expected to be high given the poverty, poor hygiene, hot/humid tropical climate and lack of access to potable water. Establishing baseline prevalence and elucidating potentially modifiable risk factors for IPI and anaemia would help public health planners, policy makers and implementers to plan and design appropriate intervention strategies to reduce associated morbidity and mortality among pre-school children.

## Materials and Methods

### Study area, design, study population, setting and period

Study design, study population and data collection sections have been previously described elsewhere (45).

### Sample size and sampling technique

The sample size of the study was determined using a single population proportion formula, considering an estimate of 24.3% expected prevalence of IPIs among children younger than 5 years old (35). Assuming any particular outcome to be within a 5% marginal error and a 95% confidence interval of certainty, the final sample size with a design effect of two and a 90% response rate was determined to be 610 mothers/children pair.

We used a multistage stratified sampling technique to identify study participants after the kebelles ((a kebelle is the smallest local government administrative unit in Ethiopia) were stratified. In the selected kebelles, 2,674 children aged 6–59 months were identified with their respective households using the registration at health posts and through the health extension workers (HEW).

We allocated the calculated sample proportionally to the selected kebelles based on the total number of households with children aged 6–59 months in each kebelle. Study participants were then identified using simple random sampling of the households. In cases where households had more than one eligible child, the eldest child was included. Accordingly, the distribution of households with respect to the kebelles was, 133 from Tsawnet, 142 from Harena, 158 from Serawat, and 177 from Mynebri.

### Data collection

After written consent was obtained from mothers or guardians of eligible children, socioeconomic, environmental, behavioural and health related data were collected using a structured questionnaire (translated from English and printed in the local Tigrigna language). Data were collected using a face-to-face administrated questionnaire and an observation method by trained data collectors, under the supervision of the principal investigators. Child hand cleanness and nail status in addition to toilet availability were assessed by direct observation.

### Variables Dependent variables

IPIs and anaemia among children aged 6-59 months (pre-school children).

### Independent variables

#### Socio-economic variables

Gender and age of the study child, mother’s/guardian’s educational status and occupation, family size, family income, number of children 6-59 months in the household.

#### Environmental and Behavioural variables

Consumption of raw vegetables, child contact with pet animals, child deworming, habit of playing in soil, shoe wearing habit, child hand cleanliness and fingernail status. Use of soap for hand washing, water source, use of household water treatment, latrine availability, and type of drinking water source.

### Faecal sample collection

Following the completion of consent and questionnaire, a clean, wide screw capped plastic stool cup, labelled with names was provided to each mother/ guardian. They were requested to bring about 10 g (thumb size) fresh stool from their child to the nearby health posts the following day within 10-30 min of passage. Participants who were not able to provide a sample on the first day were asked again on the following day.

### Parasitological analysis

Stool specimens were analysed at the respective health posts by three trained laboratory technicians using direct saline wet mount, formalin ethyl acetate concentration technique (46), and single Kato–Katz technique (thick smear 41.7 mg) (47). For the detection of *Cryptosporidium* spp. oocysts, modified Ziehl-Neelsen (MZN) staining was performed (48). Kato–Katz, wet mount preparations and modified Ziehl-Neelsen were analysed within 1h of preparation in each respective health posts to detect hookworm eggs, protozoa parasites (*E. histolytica/dispar* and *G. lamblia*), and *Cryptosporidium* spp., respectively.

The remaining stool specimens were transported in screw-capped cups in 10 % formalin to Mekelle University Medical Microbiology Laboratory and were examined using the concentration method within 8 hours after collection. After 72 hours, Kato–Katz preparations were re-examined to detect helminth ova. A child was categorised as infected if the stool sample was positive for any parasite by any of the methods used. To ensure quality, each slide was examined twice by two of the three experienced laboratory technician independently.

### Quality control

To control our data quality, 10% of the total positive specimens were randomly selected and re-examined by three experienced laboratory technicians who did not have any information about the previous results. The results of the new laboratory examinations were therefore used as quality control.

### Haemoglobin (Hb) measurement

Finger-prick blood specimens were obtained from participants to assess Hb levels using a HemoCue analyser in the health post (HemoCue Hb 201z, Sweden) (49). The apparatus was calibrated daily using the reference micro cuvettes as indicated by the manufacturer. Definition and classification of anaemia were according to the WHO cut-offs (50).

### Data analysis

Data were analysed with Stata Release 15. Confidence intervals for prevalence were calculated using the Agresti-Coull formula. Negative binomial regression was used to model **prevalence rate ratios**. Prevalence rate ratios have several advantages over odds ratios. The first is that they are simple to interpret - they directly compare prevalence, so a prevalence ratio of 2 means that prevalence is twice as high. Second, prevalence ratios, but not odds ratios, have a mathematical property called collapsibility; this means that the size of the risk ratio will not change if adjustment is made for a variable that is not a confounder (51)(52). All reported p-values were two-tailed, and statistical values were considered significant when p < 0.05.

### Ethical Clearance

Ethical clearance was obtained from Mekelle University; College of Health Science Institutional Review Board (IRB) [ERC 0844/2016]. Permission was obtained from the Tigray Regional Health Bureau and respective district Health Bureau. Written consent was obtained from mothers / guardians. Children with IPIs and anaemia were referred to the nearby health institutions and treated from the project fund.

## Results

### Prevalence of intestinal parasitosis and anaemia

Among the 610 participating pre-school children, 354(58%) (95% CI: 54.1–62.1%) were infected by one or more parasitic organism. Single, double, and triple parasitic infections were seen in 249 (40.8%), 83(13.6%), and 22 (3.6%) children, respectively. Seven different intestinal parasitic organisms were detected with *E. histolytica/dispar* the most prevalent 220(36.1%) followed by *G. lamblia* 128 (20.1%), and *H. nana* 102(16.7%). Mixed infections were common among children positive for *G. lamblia, E. histolytica/dispar* and *Cryptosporidium* spp. The prevalence of any type of anaemia was 21.6% (95% CI: 18.5% to 25.1%) (**Table 1**).

**Table 1.** Intestinal parasite and anaemia among 610 pre-school children in the Tigray region of Northern Ethiopia, 2017

| **VARIABLES** | **No (%)** |
| --- | --- |
| Children with intestinal parasite(s) detected | 354(58) |
| Number of single parasitic infections | 249(40.8) |
| Number of double parasitic infections | 83(13.6) |
| Number of triple parasitic infections | 22(3.6) |
| **Haemoglobin level** |  |
| Normal | 478(78.4) |
| Anaemia | 132(21.6) |
| **Infection type** |  |
| **Protozoa** |  |
| *E. histolytica/dispar* | 125(20.5) |
| *G. lamblia* | 52(8.5) |
| *Cryptosporidium* spp. | 11(1.8) |
| *E. vermicularis* | 2(0.3) |
| **Helminths** |  |
| *H. nana* | 59(9.7) |
| *A. lumbricoides* | 1(0.2) |
| *S. stercoralis* | 1(.2) |
| **Double infection** |  |
| *G. lamblia* + *E. histolytica/dispar* | 44(7.2) |
| *G. lamblia* + *H. nana* | 8(1.3) |
| *E. histolytica/dispar* + *H. nana* | 27(4.7) |
| *G. lamblia* + *Cryptosporidium* spp. Oocyst | 2(0.3) |
| *E. histolytica/dispar* + *E. vermicularis* | 2(0.3) |
| **Triple infection** |  |
| *G. lamblia* + *E. histolytica/dispar* + *H. nana* | 8(1.3) |
| *E. histolytica/dispar* + *G. lamblia* + *Cryptosporidium* spp. | 14(2.3) |

Prevalence of intestinal parasitic infection rose from 50% in children aged under 2 years, rising to 66% in children aged 4 to 5 years. The prevalence rate increased by 7·8% per year over the age range studied (negative binomial regression, IRR = 1·078, p=0·009). Children with unclean hands were 63.4% times more infected with IPIs than their counter parts. Similarly, prevalence of anaemia rose from 28% in those aged 6 months to one year to 44% in children aged 1 year. Thereafter it declined sharply with age to reach 7% in those aged 4 years. Because prevalence declined from one year onwards but was lower in children aged under one year, we calculated the effects of risk factors adjusting for age coded as two variables: age in years, and a binary variable indicating that the child was aged under one year (**Table 2**).

**Table 2.**
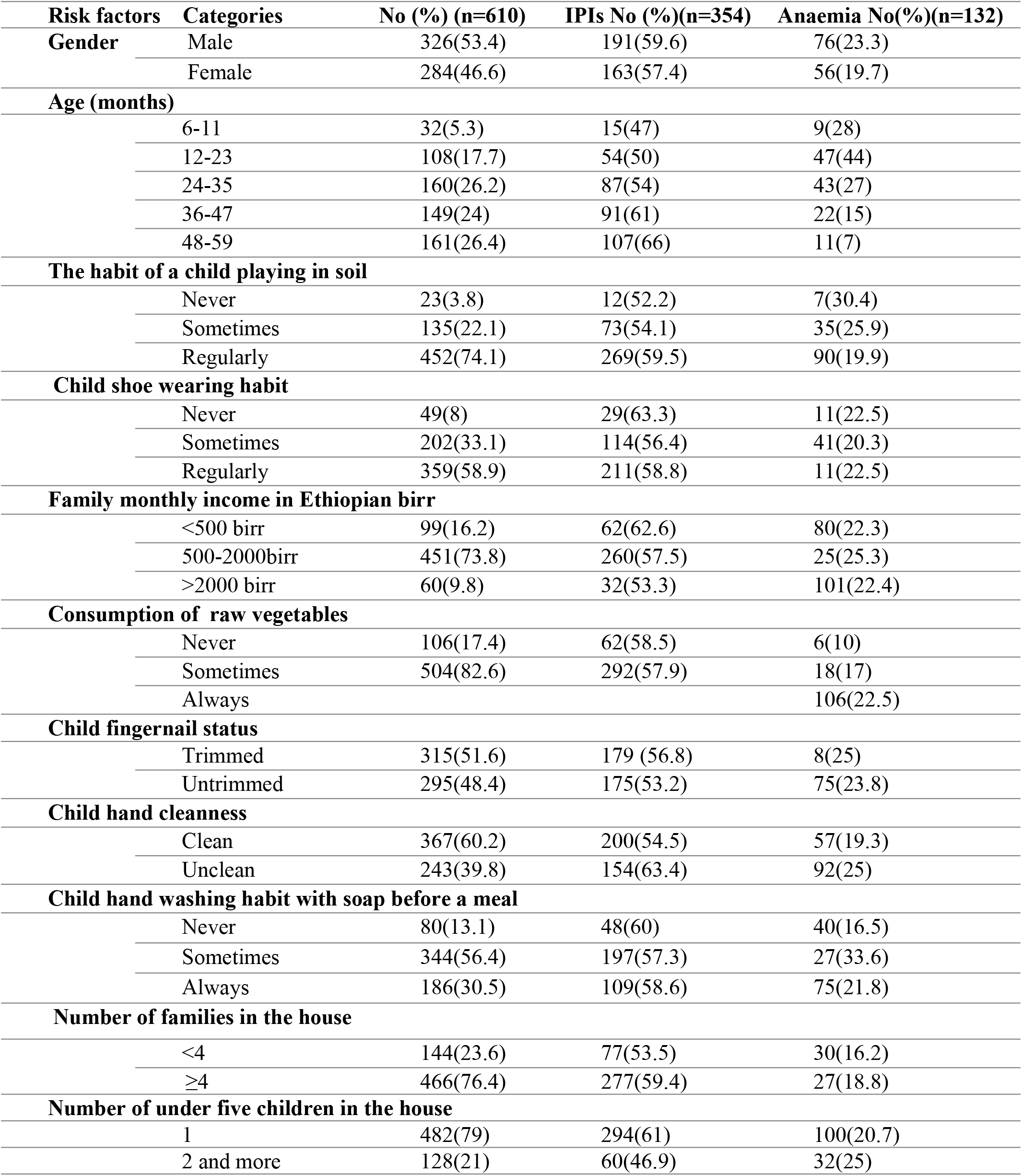
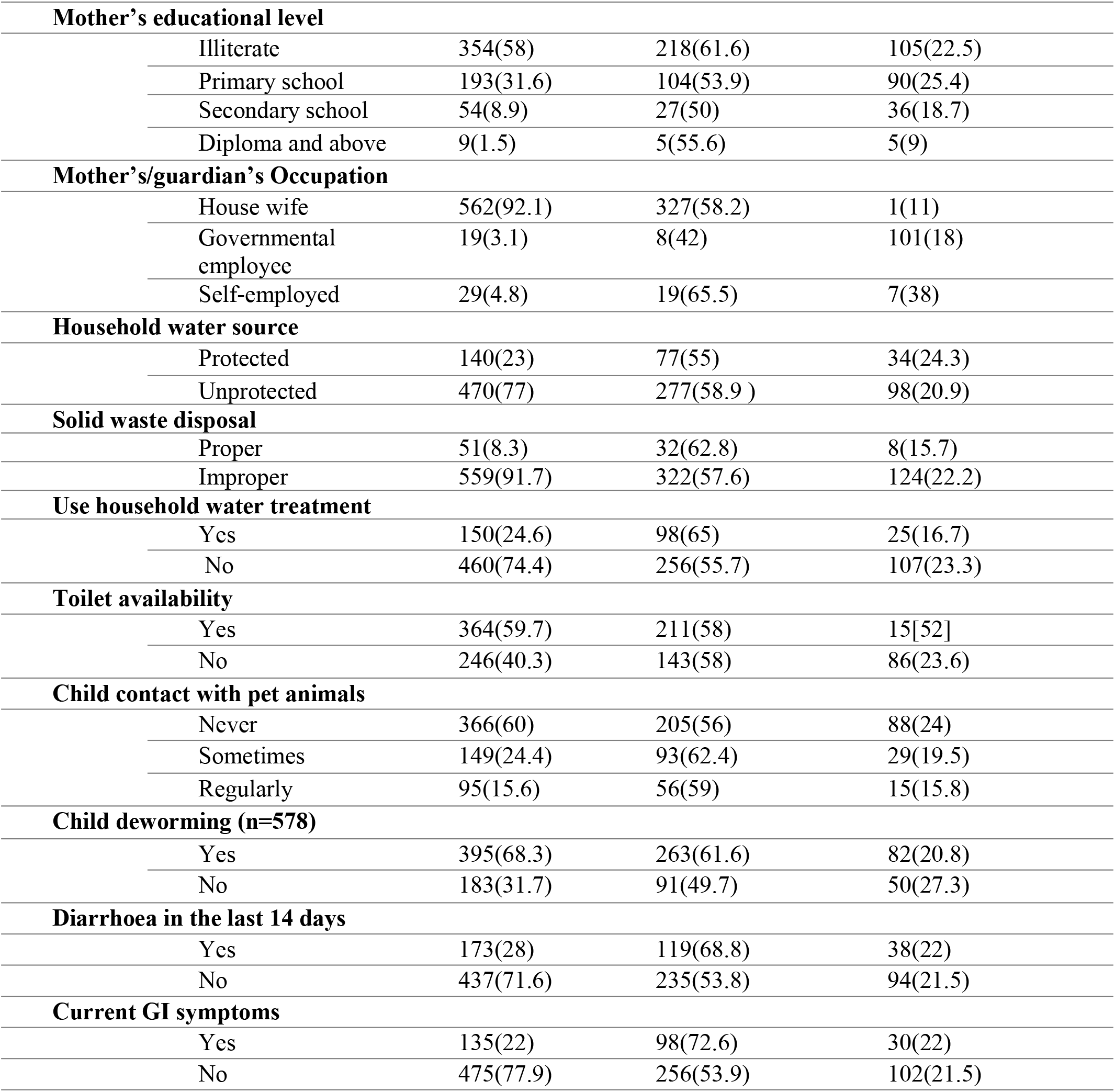
Distribution of IPIs and Anaemia among 610 pre-school children in Tigray, Ethiopia, 2017.

### Factors associated with IPIs among pre-school age children

Because prevalence of parasites is associated with age, we used negative binomial regression adjusted for age to calculate prevalence rate ratios associated with each of the associated risk factors. Adjusted for age, prevalence was higher in children with current GI symptoms (p= **0.001),** who had diarrhea in the previous 14 days (p= **0.000**), and who had been de-wormed (p= **0.045**). Prevalence was however, lower in households with two or more children aged under 5 (p=**0.015**) (**Table** 3). In multivariable analysis, having two or more children aged under five and child deworming remained significantly associated with parasite prevalence (p< 0.05).

**Table 3.** Associated Risk factors for prevalence of IPIs among 610 pre-school children in the Tigray, Ethiopia, 2017.

| Risk factor | Prevalence ratio | 95% CI | p value |
| --- | --- | --- | --- |
| 2+ under five children in the house | 0.76 | 0.61–0.95 | 0.015** |
| Any education | 0.89 | 0.74–1.10 | 0.186 |
| Child's hands clean | 0.89 | 1.0–0.80 | 0.129 |
| Solid waste disposed of properly | 0.91 | 0.76–1.10 | 0.327 |
| 4 or more children | 0.93 | 0.72–1.20 | 0.611 |
| Water from improved source | 1.0 | 0.85–1.10 | 0.382 |
| Child's fingernails trimmed? | 1.0 | 0.89–1.10 | 0.588 |
| Eats fruit or veg raw | 1.0 | 0.82–1.20 | 0.798 |
| Presence of domestic animals in house | 1.0 | 0.78–1.30 | 0.939 |
| Soap used in handwashing | 1.0 | 0.89–1.10 | 0.988 |
| Latrine availability in the house | 1.0 | 0.92–1.10 | 0.949 |
| Male sex | 1.0 | 0.89–1.20 | 0.714 |
| Habit of raw meat eating of the child | 1.0 | 0.75–1.40 | 0.855 |
| Child in contact with animals | 1.1 | 1.0–1.20 | 0.230 |
| Any skin disease | 1.1 | 0.95–1.20 | 0.258 |
| House hold water treatment | 1.2 | 1.0–1.40 | 0.057 |
| Did the child get dewormed* | 1.2 | 1.0–1.40 | 0.045** |
| Diarrhea in the last 14 days | 1.3 | 1.2–1.40 | 0.000** |
| Current GI symptoms | 1.3 | 1.1–1.60 | 0.001** |

\*\*= statistically significant, \*=this item is labelled "for above 2 years, but there are data for all children. All prevalence ratios are adjusted for age using negative binomial regression

### Factors associated with anaemia among pre-school age children

Prevalence of anaemia rose from 28% in those aged 6 months to one year to 44% in children aged 1 year. Thereafter it declined sharply with age to reach 7% in those aged 4 years. Because prevalence declined from one year onwards but was lower in children aged under one year, we calculated the effects of risk factors adjusting for age coded as two variables: age in years, and a binary variable indicating that the child was aged under one year. Eating raw meat (p= 0.000), any maternal education (p= 0.000), and household water treatment (p= 0.044), were associated with lower prevalence of anaemia, when adjusted for age. On the other hand, presence of domestic animals in the house (p=0.006), and proper disposal of solid waste (p=0.009) were both associated with increased risk (Table 4).

**Table 4.** Risk factors for anaemia among 610 pre-school children in the Tigray, Ethiopia, 2017.

| Risk factors | Prevalence ratio | 95% CI | P- value |
| --- | --- | --- | --- |
| Habit of raw meat eating of the child | 0.49 | 0.45–0.54 | 0.000** |
| 4 or more children | 0.64 | 0.39–1.00 | 0.075 |
| Any maternal education | 0.64 | 0.52–0.79 | 0.000** |
| House hold water treatment | 0.75 | 0.56–1.00 | 0.044** |
| Soap used in handwashing | 0.85 | 0.64–1.10 | 0.279 |
| Child in contact with animals | 0.87 | 0.6–1.20 | 0.399 |
| Any skin disease | 0.95 | 0.56–1.60 | 0.839 |
| Diarrhea in the last 14 days | 1.0 | 0.66–1.40 | 0.832 |
| Current GI symptoms | 1.0 | 0.59–1.60 | 0.898 |
| Did the child get dewormed* | 1.0 | 0.86–1.20 | 0.891 |
| Water from improved source | 1.0 | 0.82–1.30 | 0.752 |
| Child's fingernails trimmed | 1.1 | 1.0–1.20 | 0.077 |
| Male sex | 1.1 | 1.0–1.30 | 0.134 |
| Child's hands clean | 1.2 | 0.87–1.60 | 0.280 |
| Latrine availability in the house | 1.2 | 0.84–1.70 | 0.308 |
| Number of under-five children in the house | 1.3 | 0.92–1.90 | 0.129 |
| Eats fruit or veg raw | 1.4 | 0.91–2.00 | 0.133 |
| Presence of domestic animals in house | 1.5 | 1.1–2.00 | 0.006** |
| Solid waste disposed of properly | 1.5 | 1.1–2.10 | 0.009** |

\*\*= statistically significant, \*This item is labelled "for above 2 years, but there are data for all children. All prevalence ratios are adjusted for age using negative binomial regression

## Discussion

Locally relevant epidemiological data on IPI and anaemia prevalence among pre-school children in rural communities and identification of potentially modifiable predisposing factors, is essential to design appropriate intervention strategies. We report an overall prevalence of IPIs and anaemia of 58 % and 21.6%, respectively and have identified inadequate sanitation and clean water supplies and a requirement for family hand hygiene education as potentially modifiable associated factors. Improvements in these areas could address both short and long-term consequences of these conditions in this vulnerable population.

IPI prevalence (58 %) in our study was comparable with previous reports from Karachi, Pakistan 52.8% (53), and Benue, Nigeria 51.4% (54), though lower than studies conducted in Wondepgent, South Ethiopia 85.1% (40), Matanzas, Cuba 71.1%(55), and West Malaysia 76.5% (56). However, our prevalence is higher than previous similar studies conducted in other parts of Ethiopia such as: Wonji, 24.3% (35), Butajira, South Ethiopia 4.9% (57); Gonder, North West Ethiopia 17.3% (39), Dessie 15.5% (58), Jimma, West Ethiopia 49.6% (59), Addis Ababa 27.5% (60), Shoa, Ethiopia 17.4% (61), Dembiya, Northwest Ethiopia 25.8% (62), Hawassa, South Ethiopia 26.6 % (63), and Yirgalem Hospital, South Ethiopia 36.52% (37). Others have reported lower IPI prevalence in Islesa, Nigeria 23.3% (64), Ebonyi, Nigeria 13.7% (65), Gonbad-e Karus, Iran 26.6 % (66), Infaka, Tanzania 9.1%(67), Hoima district, rural Western Uganda 26.5% (68), Riyadh, Saudi Arabia 17.7% (69), Gaza, Palestine, 16.6%(70), and Sana’a city, Yemen, 30.9% (71). These reported differences in IPI prevalence might be due to the difference in parasitological methods used, geographical location, level of environmental sanitation, drinking water source, season, family education, personal hygiene, parental socioeconomic, and cultural difference of the study participants.

One explanation for the high IPI prevalence in this study could be because of seasonal variation. Data collection for our study took place during the rainy season in Ethiopia, other studies [35, 52, 54, 56–58, 61, 64, 69] were conducted in the dry season. Seasonal variation may be explained by increased contamination of water sources (e.g., rivers, streams, and wells) with human excreta from open defecation which is the main risk factor for diarrhoeal disease and IPIs, especially children who routinely play in the an unhygienic environment (45). In addition, as 77% of households in our study used unprotected water sources, the main factor for faecal-oral transmitted disease such as IPIs, this may also contribute to the high IPI prevalence.

Another possible reason for the high prevalence of IPIs in our study is the laboratory methodology we employed. We used the modified Ziehl Nelson method to detect *Cryptosporidium* spp, whereas the other studies except [37] and [66] did not. Furthermore, some studies isolated only the soil-transmitted helminths(STH) and not protozoa which would artificially decrease IPI prevalence(57)(35)(68) (56).

*E. histolytica, G. lambilia* and *C. parvum* were the most prevalent protozoan parasites in our study cohort. The high prevalence of *E. histolytica/dispar* (36.1%) in our study is in agreement with previous reports (54,58,72). Consequences of childhood *E. histolytica* infection include malnourishment, anaemia and stunted growth (25). However, others have reported *G. lamblia* (55,69–71,73,74), and *C. parvum* (67) as the dominant parasites. The difference in prevalence of enteric protozoa may be due to differences in contamination of drinking water sources, availability of toilets, handwashing and consumption of raw vegetables of the study participants.

Helminthic intestinal parasites, particularly soil transmitted helminths (STHs), commonly infect children in low to middle-income countries. In our study, helminthic infections were identified in 104 (17.1 %) of the children with *H. nana* 102 (16.7%) being dominant while *Ascaris*, hookworm and *Trichuris* infections were very rare or absent. The low STH incidence in our study might be due to the initiation of a national deworming program, as the majority, 68.3% of the children were dewormed during the data collection time and deworming is given on a regular basis. The relatively high prevalence of wearing shoes either regularly or sometimes and low prevalence of consumption of raw vegetables may also partially explain the low STH prevalence. In addition, the prevalence of STH might also be due to differences in environmental factors such as climate, topography (75), surface temperature, altitude, soil type and rainfall (76).

Our study shows that dewormed children were more infected with IPIs than their counter part which is against the established fact. This could be due to the fact that the prevalence of STHs, where deworming is given for, in our study was very small, and the protozoan infection which deworming has less effective against, was the most dominant. A lower prevalence of STH compared with *H. nana* has been previously reported in Peru (77) and in Eastern Ethiopia in elementary school children receiving regular albendazole deworming treatment (78). This could be probably due to the fact that albendazole has little effect on *H. nana* unlike *Ascaris lumbricoides, Trichuris trichiura* and hookworm (79).

Children aged 48-59 months (PR=1.078, p=0.009) were more likely to be infected by IPIs compared with younger children. This likely reflects increased risk of IPI exposure from playing activity of the older, more mobile children within unhygienic external environments and hence exposed to faecal-contaminated soil.

Only a quarter of households used household water treatments, mainly chlorination and boiling. Though not statistically significant, children whose family did not treat their water were 1.2 times (p= 0.057) more likely to be infected by IPI. Chlorination is less expensive, less time consuming and provides residual disinfection to protect against recontamination and significantly reduces the risk of diarrhoea (80). However, few families in our study performed chlorination because of the associated smell and taste and informed the investigators that their children prefer to drink untreated water instead. This was consistent with reports from elsewhere (81) (82). Likewise, sufficient heating, consistent and correct boiling of water reduces the childhood diarrhoea. Nevertheless, this increases fuel consumption, is time consuming, and because of lack of residual protection, treated water can be re-contaminated following the immersion of unclean fingers, other fomites, or through storage in a dirty container (83). Hence a cheap, point of use technology which overcomes these limitation is required by such communities to tackle the consumption of unsafe drinking water.

The prevalence of childhood anaemia among pre-school children in our study was 21.6%, which is comparable with studies from South central Ethiopia 28.2% (84), Western province, Kenya 25% (85), West Malaysia 26.2% (56), and Northeast Brazil 28.9% (86). Our study was, however, lower than other reports from Ethiopia 37.3% −42.2% (43) (87), the 2016 Ethiopian Health Demographic Survey, 57 % [33], the EDHS anaemia prevalence of the Tigray regional state, 54% [33] and the WHO classification for anaemia 40% [18]. Likewise, studies performed outside Ethiopia have reported higher anaemia prevalence including those conducted in Pernambuco, Brazil 32.8% (88), North Western Uganda 37.2% (21), Uganda 72% and 60% (23), Edo state, Nigeria 38.6% (89), Bangladesh 51.9% (90), Senegal 53.4% (91), Arusha District, Tanzania 84.6% (17), Cape Verde, West Africa 51.8% (92), Simanjiro district, Tanzania 47.6% (93), Indonesia 56.1% (94), Namutumba district, Uganda 58.8% (95), sub-Saharan Africa 59.9% (96), and Brazil 45.1% (88).

Possible reasons for the variation in anaemia prevalence could be due to differences in maternal education [84,69,78], concomitant childhood malaria superinfection [21–23,77,81], family income [21,23,43,69,76,81,85], drinking water source [78, 81], personal hygiene (97), and STH infections [23,69,80,84]. For example, the high anaemia rate in the studies such as Arusha District, Tanzania, Kenya, Senegal, Uganda, South central Ethiopia, was due to concomitant malaria infection. Seasonality may have also affected the anaemia prevalence in our study as malaria prevalence is usually low during the rainy season of the country (June-August). In addition, our study participants were from the community rather than from health institutions, and the prevalence of STH as the main cause of anaemia [23, 69, 80, 84,(98) was very low in our study. Our multivariate logistic regression analysis shows that children aged 12-23 months were more likely to have anaemia compared with younger age groups. This association of age and anaemia has been previously reported [43, 76, 81, 83].

## Limitation of the study

One of the limitations of our study was that differentiation between the morphologically identical species of *Entamoeba* was not within the scope of this study, as only conventional microscopy was used to detect the amoebae.

## Conclusions

This study revealed the significant burden of IPI and anaemia in preschool children in a rural community in Northern Ethiopia. More than half of the children were infected with intestinal parasites and one in five were anaemic. Age of the child, child deworming and having two or more children aged under five were significant predictors for the IPIs in the study areas. More investment and commitment to addressing the problem of childhood IPIs by designing and implementing prevention strategies, such as integrating mothers’/guardian’s education on personal and environmental hygiene into existing national health extension program.

While anemia was comparably lower, it was associated with age, mother’s/guardians educational status, and household water treatment. Given the identified burden of childhood IPI and anaemia during the period of life most critical for physical and cognitive development (i.e., the preschool years), this study underlines the need for interventions focusing on the identified modifiable risk factors to prevent long-term morbidity and give these children the maximal opportunity for health in the future.

## Acknowledgments

We would like to thank all the children, parents, guardians and caregivers for their collaboration. The cooperation of the Tigray Regional Health Bureau and respective Health bureaus of the districts are also highly acknowledged by the authors. The authors also like to express their sincere gratitude to the health extension workers in each site for their help in data collection and communication with each mother/guardian at a household level.

## Author Contributions

AGW, TA, MT, LN, DY, KMG, JM designed the study. AGW, HTA, RC worked on the analysis and interpretation of the data. AGW, prepared the manuscript for publication. AGW, TA, MT, HTA, JM, FF, KMG, RC reviewed the manuscript for publication. All authors read and approved the final paper.

## Legend 1. Supporting Information Legends

Title: Raw data working

## References

1. Lobo ML, Augusto J, Antunes F, Ceita J, Xiao L, Codices V, et al. Cryptosporidium spp., Giardia duodenalis, Enterocytozoon bieneusi and other intestinal parasites in young children in Lobata Province, Democratic Republic of São Tomé and Principe. PLoS One. 2014;9(5).

2. Hotez PJ, Fenwick A, Savioli L, Molyneux DH. Rescuing the bottom billion through control of neglected tropical diseases. Lancet. 2009;373(9674):1570–5.

3. Keiser J, Utzinger J. The Drugs We Have and the Drugs We Need Against Major Helminth Infections. Adv Parasitol. 2010;73(C):197–230.

4. Fischer Walker CL, Aryee MJ, Boschi-Pinto C, Black RE. Estimating diarrhea mortality among young children in low and middle income countries. Vol. 7, PLoS ONE. 2012.

5. Keogh MB, Castro-Alférez M, Polo-López MI, Fernández Calderero I, Al-Eryani YA, Joseph-Titus C, et al. Capability of 19-L polycarbonate plastic water cooler containers for efficient solar water disinfection (SODIS): Field case studies in India, Bahrain and Spain. Sol Energy. 2015;116(June 2015):1–11.

6. Eckmann L. Mucosal defences against Giardia. Vol. 25, Parasite Immunology. 2003. p. 259–70.

7. Yaoyu F, Xiao L. Zoonotic potential and molecular epidemiology of Giardia species and giardiasis. Clin Microbiol Rev. 2011;24(1):110–40.

8. Xiao L. Molecular epidemiology of cryptosporidiosis: An update. Exp Parasitol. 2010;124(1):80–9.

9. Chalmers RM, Davies AP. Minireview: Clinical cryptosporidiosis. Vol. 124, Experimental Parasitology. 2010. p. 138–46.

10. World Health Organization. Soil-transmitted helminth infections [Internet]. WHO Fact Sheet. 2018. Available from: http://www.who.int/news-room/fact-sheets/detail/soil-transmitted-helminth-infections

11. Global Burden of Disease. Estimates of global, regional, and national morbidity, mortality, and aetiologies of diarrhoeal diseases: a systematic analysis for the Global Burden of Disease Study 2015. Lancet Infect Dis. 2017;17(9):909–48.

12. Thomas Iv LJ, Zweig AP, Tosh AK. An adolescent with chronic giardiasis mimicking anorexia nervosa. Int J Adolesc Med Health. 2014;26(2):293–5.

13. Al-Mekhlafi HM, Al-Maktari MT, Jani R, Ahmed A, Anuar TS, Moktar N, et al. Burden of Giardia duodenalis Infection and Its Adverse Effects on Growth of Schoolchildren in Rural Malaysia. PLoS Negl Trop Dis. 2013;7(10).

14. Nematian J, Gholamrezanezhad A, Nematian E. Giardiasis and other intestinal parasitic infections in relation to anthropometric indicators of malnutrition: a large, populationbased survey of schoolchildren in Tehran. Ann Trop Med Parasitol [Internet]. 2008;102(3):209–14. Available from: http://www.tandfonline.com/doi/full/10.1179/136485908X267876

15. Hall A, Hewitt G, Tuffrey V, De Silva N. A review and meta-analysis of the impact of intestinal worms on child growth and nutrition. MCN Matern Child Nutr. 2008;4:118–236.

16. McLean E, Cogswell M, Egli I, Wojdyla D, de Benoist B. Worldwide prevalence of anaemia, WHO Vitamin and Mineral Nutrition Information System, 1993-2005. Public Heal Nutr [Internet]. 2009;12(4):444–54. Available from: http://whqlibdoc.who.int/publications/2008/9789241596657_eng.pdf

17. Kejo D, Petrucka PM, Martin H, Kimanya ME, Mosha TC. Prevalence and predictors of anemia among children under 5 years of age in Arusha District, Tanzania. Pediatr Heal Med Ther [Internet]. 2018;9:9–15. Available from: http://www.ncbi.nlm.nih.gov/pubmed/29443328%0Ahttp://www.pubmedcentral.nih.gov/articlerender.fcgi?artid=PMC5804135

18. World Health Organization. the Global Prevalence of Anaemia in 2011. WHO Rep [Internet]. 2011;48. Available from: http://apps.who.int/iris/bitstream/10665/177094/1/9789241564960_eng.pdf?ua=1

19. World Health Organization. the Global Prevalence of Anaemia in 2011 [Internet]. WHO Library Cataloguing-in-Publication Data. 2015. 48 p. Available from: www.who.int/about/licensing/copyright_form/en/index.html%0Ahttp://apps.who.int/iris/bitstream/10665/177094/1/9789241564960_eng.pdf?ua=1

20. Phiri KS, Calis JCJ, Faragher B, Nkhoma E, Ng’oma K, Mangochi B, et al. Long term outcome of severe anaemia in Malawian children. PLoS One [Internet]. 2008;3(8):e2903. Available from: http://www.ncbi.nlm.nih.gov/pubmed/18682797

21. Legason ID, Atiku A, Ssenyonga R, Olupot-Olupot P, Barugahare JB. Prevalence of Anaemia and Associated Risk Factors among Children in North-western Uganda: A Cross Sectional Study. BMC Hematol. BMC Hematology; 2017;17(1):1–9.

22. Ngesa O, Mwambi H. Prevalence and risk factors of anaemia among children aged between 6 months and 14 years in Kenya. PLoS One. 2014;9(11).

23. Menon MP, Yoon SS. Prevalence and Factors Associated with Anemia Among Children Under 5 Years of Age—Uganda, 2009. Am J Trop Med Hyg [Internet]. 2015;93(3):521–6. Available from: http://www.ajtmh.org/content/journals/10.4269/ajtmh.15-0102

24. Falkingham M, Abdelhamid A, Curtis P, Fairweather-Tait S, Dye L, Hooper L. The effects of oral iron supplementation on cognition in older children and adults: A systematic review and meta-analysis. Nutr J. 2010;9(1).

25. Mondal D, Petri WA, Sack RB, Kirkpatrick BD, Haque R. Entamoeba histolytica-associated diarrheal illness is negatively associated with the growth of preschool children: evidence from a prospective study. Trans R Soc Trop Med Hyg. 2006;100(11):1032–8.

26. Cassat JE, Skaar EP. Iron in infection and immunity. Vol. 13, Cell Host and Microbe. 2013. p. 509–19.

27. Oppenheimer SJ. Iron and its relation to immunity and infectious disease. J Nutr [Internet]. 2001;131(2S–2):616S–633S; discussion 633S–635S. Available from: http://www.ncbi.nlm.nih.gov/pubmed/11160594

28. More S, Shivkumar VB, Gangane N, Shende S. Effects of iron deficiency on cognitive function in school going adolescent females in rural area of central India. Anemia. 2013;2013.

29. Halterman JS, Kaczorowski JM, Aligne CA, Auinger P, Szilagyi PG. Iron Deficiency and Cognitive Achievement Among School-Aged Children and Adolescents in the United States. Pediatrics [Internet]. 2001;107(6):1381–6. Available from: http://pediatrics.aappublications.org/cgi/doi/10.1542/peds.107.6.1381

30. Soewondo S, Husaini M, Pollitt E. Effects of iron deficiency on attention and learning processes in preschool children: Bandung, Indonesia [Internet]. Vol. 50, American Journal of Clinical Nutrition. 1989. p. 664–7. Available from: http://onlinelibrary.wiley.com/o/cochrane/clcentral/articles/200/CN-00062200/frame.html

31. Albonico M, Allen H, Chitsulo L, Engels D, Gabrielli AF, Savioli L. Controlling soil-transmitted helminthiasis in pre-school-age children through preventive chemotherapy. PLoS Negl Trop Dis. 2008;2(3).

32. Bobonis GJ, Miguel E, Puri-Sharma C. Anemia and School Participation. J Hum Resour [Internet]. 2006;XLI(4):692–721. Available from: http://jhr.uwpress.org/lookup/doi/10.3368/jhr.XLI.4.692

33. Fdre CSA. Key Findings Health Survey Ethiopia. 2016; Available from: https://dhsprogram.com/pubs/pdf/SR241/SR241.pdf

34. Tegegne Y, Wondmagegn T, Worku L, Jejaw Zeleke A. Prevalence of Intestinal Parasites and Associated Factors among Pulmonary Tuberculosis Suspected Patients Attending University of Gondar Hospital, Gondar, Northwest Ethiopia. J Parasitol Res [Internet]. 2018;2018:1–6. Available from: http://www.ncbi.nlm.nih.gov/pubmed/29666698%0A http://www.pubmedcentral.nih.gov/articlerender.fcgi?artid=PMC5832163%0Ahttps://www.hindawi.com/journals/jpr/2018/9372145/

35. Ghiwot Y, Degarege A, Erko B. Prevalence of intestinal parasitic infections among children under five years of age with emphasis on schistosoma mansoni in Wonji Shoa sugar estate, ethiopia. PLoS One. 2014;9(10).

36. Wadilo F, Solomon F. Magnitude of Intestinal Parasitosis among Under Five Year Children Presenting with Acute Diarroheal Illness in South Ethiopian Hospital. 2016;31:71–7.

37. Firdu T, Abunna F, Girma M. Intestinal Protozoal Parasites in Diarrheal Children and Associated Risk Factors at Yirgalem Hospital, Ethiopia: A Case-Control Study. Int Sch Res Not [Internet]. 2014;2014:1–8. Available from: https://www.hindawi.com/archive/2014/357126/

38. Y. B, G. M, A. A, B. E, C. H, A. A, et al. Prevalence and risk factors for soil-transmitted helminth infection in mothers and their infants in Butajira, Ethiopia: a population based study. BMC Public Health [Internet]. 2010;10:21. Available from: http://ovidsp.ovid.com/ovidweb.cgi?T=JS&PAGE=reference&D=emed9&NEWS=N&AN=20085635

39. Aleka Y, G/egziabher S, Tamir W, Birhane M, Alemu A. Prevalence and Associated Risk Factors of Intestinal Parasitic Infection among Under five Children in University of Gondar Hospital, Gondar, Northwest Ethiopia. Biomed Res Ther [Internet]. 2015;2(8):20. Available from: http://www.globalsciencejournals.com/article/10.7603/s40730-015-0020-2

40. Nyantekyi LA, Legesse M, Belay M, Tadesse K, Manaye K, Macias C, et al. Intestinal parasitic infections among under-five children and maternal awareness about the infections in Shesha Kekele, Wondo Genet, Southern Ethiopia. Ethiop J Heal Dev. 2010;24(3):185–90.

41. Gari T, Loha E, Deressa W, Solomon T, Atsbeha H, Assegid M, et al. Anaemia among children in a drought affected community in south-central Ethiopia. PLoS One. 2017;12(3):1–16.

42. Deribew A, Alemseged F, Tessema F, Sena L, Birhanu Z, Zeynudin A, et al. Malaria and under-nutrition: A community based study among under-five children at risk of malaria, South-West Ethiopia. PLoS One. 2010;5(5).

43. Gebreegziabiher G, Etana B, Niggusie D. Determinants of Anemia among Children Aged 6-59 Months Living in Kilte Awulaelo Woreda, Northern Ethiopia. Anemia. 2015;2014.

44. Kotloff KL, Nataro JP, Blackwelder WC, Nasrin D, Farag TH, Panchalingam S, et al. Burden and aetiology of diarrhoeal disease in infants and young children in developing countries (the Global Enteric Multicenter Study, GEMS): a prospective, case-control study. Lancet. 2013;382:209–22.

45. Gebreyesus A, Id W, Dejene TA, Teferi M, Negash L, Yemane D, et al. Risk factors for diarrhoea and malnutrition among children under the age of 5 years in the Tigray Region of Northern Ethiopia. 2018;32–9.

46. Zeibig EA. Clinical Parasitology: A PRATICAL APPROACH. Vol. 1, Elsevier. 1997. 385 p.

47. World Health Organization. Bench aids for the diagnosis of intestinal parasites. Programme on Intestinal Parasitic Infections, Division of Communicable Diseases. 1994. 23 p.

48. Pohlenz, S.A. and Henriksen JFL. Staining of Cryptosporidia by a modified ziehl-neelsen technique. Acta vet scand. 1981;22:594–6.

49. Neufeld L, García-Guerra A, Sánchez-Francia D, Newton-Sánchez O, Ramírez-Villalobos MD, Rivera-Dommarco J. Hemoglobin measured by Hemocue and a reference method in venous and capillary blood: A validation study. Salud Publica Mex. 2002;44(3):219–27.

50. World Health Organization. Iron Deficiency Anaemia: Assessment, Prevention and Control, A guide for program managers. Control [Internet]. 2001;114. Available from: http://www.who.int/nutrition/publications/en/ida_assessment_prevention_control.pdf

51. When R, Are O. The Relative Merits of Risk Ratios and Odds Ratios. 2019;163(5):438–45.

52. Tamhane AR, Westfall AO, Burkholder GA, Cutter GR. HHS Public Access. 2017;35(30):5730–5.

53. Mehraj V, Hatcher J, Akhtar S, Rafique G, Beg MA. Prevalence and factors associated with intestinal parasitic infection among children in an urban slum of Karachi. PLoS One. 2008;3(11).

54. Tyoalumun K, Abubakar S, Christopher N. Prevalence of Intestinal Parasitic Infections and their Association with Nutritional Status of Rural and Urban Pre-School Children in Benue State, Nigeria. Int J MCH AIDS [Internet]. 2016;5(2):146–52. Available from: https://www.ncbi.nlm.nih.gov/pmc/articles/PMC5187646/pdf/IJMA-5-146.pdf

55. Cañete R, Díaz MM, Avalos García R, Laúd Martinez PM, Manuel Ponce F. Intestinal Parasites in Children from a Day Care Centre in Matanzas City, Cuba. PLoS One. 2012;7(12):1–4.

56. Ngui R, Lim YAL, Kin LC, Chuen CS, Jaffar S. Association between anaemia, iron deficiency anaemia, neglected parasitic infections and socioeconomic factors in rural children of West Malaysia. PLoS Negl Trop Dis. 2012;6(3):1–8.

57. Shumbej T, Belay T, Mekonnen Z, Tefera T, Zemene E, Ferron ES. Soil-transmitted helminths and associated factors among pre-school children in Butajira Town, southcentral Ethiopia: A community-based cross-sectional study. PLoS One. 2015;10(8).

58. Gebretsadik D, Metaferia Y, Seid A, Fenta GM, Gedefie A. Prevalence of intestinal parasitic infection among children under 5 years of age at Dessie Referral Hospital: cross sectional study. BMC Res Notes [Internet]. BioMed Central; 2018;11(1):771. Available from: https://bmcresnotes.biomedcentral.com/articles/10.1186/s13104-018-3888-2

59. Beyene G, Tasew H. Prevalence of intestinal parasite, Shigella and Salmonella species among diarrheal children in Jimma health center, Jimma southwest Ethiopia: A cross sectional study. Ann Clin Microbiol Antimicrob [Internet]. Annals of Clinical Microbiology and Antimicrobials; 2014;13(1):1–7. Available from: Annals of Clinical Microbiology and Antimicrobials

60. Adamu H, Endeshaw T, Teka T, Kifle A, Petros B. the Prevalence of Intestinal Protozoa. Lancet. 1917;189(4878):307.

61. Zemene T, Shiferaw MB. Prevalence of intestinal parasitic infections in children under the age of 5 years attending the Debre Birhan referral hospital, North Shoa, Ethiopia. BMC Res Notes [Internet]. BioMed Central; 2018;11(1):1–6. Available from: https://doi.org/10.1186/s13104-018-3166-3

62. Gizaw Z, Adane T, Azanaw J, Addisu A, Haile D. Childhood intestinal parasitic infection and sanitation predictors in rural Dembiya, northwest Ethiopia. Environ Health Prev Med. Environmental Health and Preventive Medicine; 2018;23(1):1–10.

63. Mulatu G, Zeynudin A, Zemene E, Debalke S, Beyene G. Intestinal parasitic infections among children under five years of age presenting with diarrhoeal diseases to two public health facilities in Hawassa, South Ethiopia. Infect Dis Poverty. 2015;4(1).

64. Tinuade O, John O, Saheed O, Oyeku O, Fidelis N, Olabisi D. Parasitic etiology of childhood diarrhea. Indian J Pediatr [Internet]. 2006;73(12):1081–4. Available from: http://www.scopus.com/inward/record.url?eid=2-s2.0-33846951479&partnerID=40&md5=e36e25f0ca8d911763436a3a4a2de60b

65. Achi EC, Njoku OO, Nnachi AU, Efunshile AM, Mbah JO, Aghanya IN, et al. Prevalence of intestinal parasitic infections among under five children in Abakaliki Local Government Area of Ebonyi State. Eur J Pharm Med Res. 2017;4(4):218–22.

66. Mesgarian F, Sofizadeh A, Shoraka HR, Rahimi HR, Hesari A, Soheili N, et al. Prevalence of Intestinal Parasite Infections among Children in the Day Care Centers of Gonbad-e Kavus County, North-Eastern Iran. 2017;19(10):1–6.

67. Vargas M, Gascón J, Casals C, Schellenberg D, Urassa H, Kahigwa E, et al. Etiology of diarrhea in children less than five years of age in Ifakara, Tanzania. Am J Trop Med Hyg. 2004;70(5):536–9.

68. Ojja S, Kisaka S, Ediau M, Tuhebwe D, Kisakye AN, Halage AA, et al. Prevalence, intensity and factors associated with soil-transmitted helminths infections among preschool-age children in Hoima district, rural western Uganda. BMC Infect Dis. BMC Infectious Diseases; 2018;18(1):1–12.

69. Wafa A.I and AL Megrin. Assessment the prevalence of intestinal parasitic and associated risk factors among preschool children in Riyadh, Saudi Arabia. Vol. 10, Research journal of parasitology. 2015. p. 31–41.

70. Al-hindi AI, El-kichaoi A. Occurrence of Gastrointestinal Parasites Among Pre-School. 2008;16(1):125–30.

71. Alyousefi NA, Mahdy MAK, Mahmud R, Lim YAL. Factors associated with high prevalence of intestinal protozoan infections among patients in Sana’a city, Yemen. PLoS One. 2011;6(7).

72. Mulatu G, Zeynudin A, Zemene E, Debalke S, Beyene G. Intestinal parasitic infections among children under five years of age presenting with diarrhoeal diseases to two public health facilities in Hawassa, South Ethiopia. Infect Dis Poverty [Internet]. 2015;4:49. Available from: http://www.ncbi.nlm.nih.gov/pubmed/26530964

73. S. M, H. S, T. A. Frequency and risk factors for intestinal parasitic infection in children under five years age at a tertiary care hospital in Karachi [Internet]. Vol. 59, Journal of the Pakistan Medical Association. 2009. p. 216–9. Available from: http://jpma.org.pk//PdfDownload/1667.pdf%5Cnhttp://ovidsp.ovid.com/ovidweb.cgi?T=JS&PAGE=reference&D=emed9&NEWS=N&AN=2009236721

74. Zemene T, Shiferaw MB. Prevalence of intestinal parasitic infections in children under the age of 5 years attending the Debre Birhan referral hospital, North Shoa, Ethiopia. BMC Res Notes. 2018;11(1).

75. Brooker S, Singhasivanon P, Waikagul J, Supavej S, Kojima S, Takeuchi T, et al. Mapping soil-transmitted helminths in Southeast Asia and implications for parasite control. Southeast Asian J Trop Med Public Health. 2003;34(1):24–36.

76. Appleton CC, Gouws E. The distribution of common intestinal nematodes along an altitudinal transect in KwaZulu-Natal, south africa. Ann Trop Med Parasitol. 1996;90(2):181–8.

77. Machicado JD, Marcos LA, Tello R, Canales M, Terashima A, Gotuzzo E. Diagnosis of soil-transmitted helminthiasis in an Amazonic community of Peru using multiple diagnostic techniques. Trans R Soc Trop Med Hyg. 2012;106(6):333–9.

78. Tefera E, Mohammed J, Mitiku H. Intestinal helminthic infections among elementary students of Babile town, eastern Ethiopia. Pan Afr Med J. 2015;20:1–10.

79. Horton J. Albendazole: a review of anthelmintic efficacy and safety in humans. Parasitology [Internet]. 2000;121(S1):S113. Available from: http://www.journals.cambridge.org/abstract_S0031182000007290

80. Fagerli K, Trivedi KK, Sodha S V, Blanton E, Ati A, Nguyen T, et al. HHS Public Access. 2018;145(15):3294–302.

81. Freeman MC, Quick RE, Abbott DP, Ogutu P, Rheingans R. Increasing equity of access to point-of-use water treatment products through social marketing and entrepreneurship: A case study in western Kenya. J Water Health. 2009;7(3):527–34.

82. O’Reilly CE, Freeman MC, Ravani M, Migele J, Mwaki A, Ayalo M, et al. The impact of a school-based safe water and hygiene programme on knowledge and practices of students and their parents: Nyanza Province, western Kenya, 2006. Epidemiol Infect. 2008;136(1):80–91.

83. Sodha S V., Menon M, Trivedi K, Ati A, Figueroa ME, Ainslie R, et al. Microbiologic effectiveness of boiling and safe water storage in South Sulawesi, Indonesia. J Water Health. 2011;9(3):577–85.

84. Gari T, Loha E, Deressa W, Solomon T, Atsbeha H, Assegid M, et al. Anaemia among children in a drought affected community in south-central Ethiopia. PLoS One. 2017;12(3).

85. Kisiangani I, Mbakaya C, Makokha A. Prevalence of Anaemia and Associated Factors Among Preschool Children (6-59 Months) in Western Province, Kenya. Public Heal Prev Med. 2015;1(1):28–32.

86. Carvalho AGC, de Lira PIC, Barros M de FA, Aléssio MLM, Lima M de C, Carbonneau MA, et al. Diagnóstico de anemia por deficiência de ferro em crianças do Nordeste do Brasil. Rev Saude Publica. 2010;44(3):513–9.

87. Kawo KN, Asfaw ZG, Yohannes N. Multilevel Analysis of Determinants of Anemia Prevalence among Children Aged 6-59 Months in Ethiopia: Classical and Bayesian Approaches. Anemia. 2018;2018.

88. Luciana PL, Filho MB, Israel Pedro LC De, Osório MM, I MBF, Israel P, et al. Prevalence of anemia and associated factors in children aged 6-59 months in Pernambuco, Northeastern Brazil. Rev Saúde Pública [Internet]. 2011;45(3):1–8. Available from: http://www.scielosp.org/pdf/rsp/v45n3/en_2433.pdf

89. Osazuwa F, Ayo OM, Imade P. A significant association between intestinal helminth infection and anaemia burden in children in rural communities of Edo state, Nigeria. N Am J Med Sci. 2011;3(1):30–4.

90. Khan JR, Awan N, Misu F. Determinants of anemia among 6-59 months aged children in Bangladesh: Evidence from nationally representative data. BMC Pediatr [Internet]. BMC Pediatrics; 2016;16(1):1–12. Available from: http://dx.doi.org/10.1186/s12887-015-0536-z

91. R.C. T, B. F, J.L. N, C.T. N, P. M, I.C. B, et al. Prevalence of intestinal parasites, anaemia and anthropometric status among children under five years of age in Lamarame (Senegal) [Internet]. Vol. 83, American Journal of Tropical Medicine and Hygiene. 2010. p. 93. Available from: http://ovidsp.ovid.com/ovidweb.cgi?T=JS&PAGE=reference&D=emed9&NEWS=N&AN=70442320

92. Semedo RML, Santos MMAS, Baião MR, Luiz RR, Da Veiga G V. Prevalence of Anaemia and Associated Factors among Children below Five Years of Age in Cape Verde, West Africa. J Heal Popul Nutr. 2014;32(4):646–57.

93. Nyaruhucha CN, Mamiro PS, Kerengi AJ. Prevalence of anaemia and parasitic infections among underfive children in Simanjiro District, Tanzania. Tanzan Health Res Bull [Internet]. 2005;7(1):35–9. Available from: http://proxy.lib.umich.edu/login?url= http://search.ebscohost.com/login.aspx?direct=true&db=lhh&AN=20073067359&site=ehost-live&scope=site%5Cnhttp://www.ajol.info/journal_index.php?jid=78%5Cnemail:nyaruhucha@suanet.ac.tz

94. Howard CT, de Pee S, Sari M, Bloem MW, Semba RD. Association of diarrhea with anemia among children under age five living in rural areas of Indonesia. J Trop Pediatr. 2007;53(4):238–44.

95. Kuziga F, Adoke Y, Wanyenze RK. Prevalence and factors associated with anaemia among children aged 6 to 59 months in Namutumba district, Uganda: A cross-sectional study. BMC Pediatr [Internet]. BMC Pediatrics; 2017;17(1):1–9. Available from: http://dx.doi.org/10.1186/s12887-017-0782-3

96. Moschovis PP, Wiens MO, Arlington L, Antsygina O, Hayden D, Dzik W, et al. Individual, maternal and household risk factors for anaemia among young children in sub-Saharan Africa: A cross-sectional study. BMJ Open. 2018;8(5):1–14.

97. Mahmud MA, Spigt M, Mulugeta Bezabih A, López Pavon I, Dinant G-J, Blanco Velasco R. Risk factors for intestinal parasitosis, anaemia, and malnutrition among school children in Ethiopia. Pathog Glob Health [Internet]. 2013;107(2):58–65. Available from: http://www.tandfonline.com/doi/full/10.1179/2047773213Y.0000000074

98. Verhagen LM, Incani RN, Franco CR, Ugarte A, Cadenas Y, Sierra Ruiz CI, et al. High Malnutrition Rate in Venezuelan Yanomami Compared to Warao Amerindians and Creoles: Significant Associations WITH Intestinal Parasites and Anemia. PLoS One. 2013;8(10):1–12.

